# A Novel Household Water Insecurity Scale: Procedures and Psychometric Analysis among Postpartum Women in Western Kenya

**DOI:** 10.1101/294298

**Authors:** Godfred O. Boateng, Shalean M. Collins, Patrick Mbullo, Pauline Wekesa, Maricianah Onono, Torsten B. Neilands, Sera L. Young

## Abstract

Our ability to measure household-level food insecurity has revealed its critical role in a range of physical, psychosocial, and health outcomes. Currently, there is no analogous, standardized instrument for quantifying household-level water insecurity, which prevents us from understanding both its prevalence and consequences. Therefore, our objectives were to develop and validate a household water insecurity scale appropriate for use in our cohort in western Kenya. We used a range of qualitative techniques to develop a preliminary set of 29 household water insecurity questions, and administered those questions at 15 and 18 months postpartum, concurrent with a suite of other survey modules. These data were complemented by data on quantity of water used and stored, and microbiological quality. Inter-item and item-total correlations were performed to reduce scale items to 20. Exploratory factor and parallel analyses were used to determine the latent factor structure; a unidimensional scale was hypothesized and tested using confirmatory factor and bifactor analyses, along with multiple statistical fit indices. Reliability was assessed using Cronbach’s alpha and the coefficient of stability, which produced a coefficient alpha of 0.97 at 15 and 18 months postpartum and a coefficient of stability of 0.62. Predictive, convergent and discriminant validity of the final household water insecurity scale were supported, based on relationships with food insecurity, perceived stress, per capita household water use, and time and money spent acquiring water. The resultant scale is a valid and reliable instrument. It can be used in this setting to test a range of hypotheses about the role of household water insecurity in numerous physical and psychosocial health outcomes, to identify the households most vulnerable to water insecurity, and to evaluate the effects of water-related interventions. To extend its applicability, we encourage efforts to develop a cross-culturally valid scale using robust qualitative and quantitative techniques.

## Introduction

Water security, the ability to access and benefit from affordable, adequate, reliable, and safe water for wellbeing and a healthy life [1], is fundamental to physical and mental health [2– 5]. There is also widespread agreement that difficulty with regular availability and access to water in sufficient quality and quantity is a serious problem [6,7] that will only increase, given climatic changes and increased water use [8,9]. While there are many plausible ways in which water insecurity can impact health, there are very few empirical data exploring the pathways by which water insecurity may be deleterious [10].

Understanding the impact of water insecurity is perhaps most pressing for two groups: women and people living with HIV (PLHIV) residing in places with low water security. In most parts of the developing world, women bear the physical responsibility and psychological burden of ensuring adequate household water [4,11–13]. This responsibility can be very demanding in terms of time and energy (e.g. walking long distances to water sources, carrying heavy jerry cans) and can also leave women vulnerable to physical and sexual violence *en route* to sources [14,15]. Further, water acquisition can leave women less time for other critical responsibilities (which are also often water-intensive and promote health and hygiene), such as bathing children, laundry, and preparing food [16]. The energy and time required to acquire water can also compromise women’s ability to care for their children through activities such as breastfeeding and clinic visits. It can preclude women from engaging in wage-earning activities and children from attending school [10,13]. Finally, pregnant and lactating women can have less physical ability to access water just as their needs increase, making the need for readily accessible, clean water especially vital during pregnancy and lactation [17].

Because water insecurity often occurs in regions of high HIV prevalence, there is a likelihood of syndemicity, the co-occurrence of intersecting, overlapping epidemics [18–20]. Food insecurity and HIV have similarly been considered as syndemic [21]. PLHIV are more susceptible to waterborne diseases, including diarrhea [22]. Diarrhea can in turn lead to poor intestinal absorption of essential nutrients and therapeutic dosages of medicines [23,24]. People with advanced HIV can have compromised physical ability to access water [25], and their care often requires more water to preserve hygiene [25–27]. Further, an additional 1.5 liters of clean water per day is required to metabolize some HIV medicines [26].

Of the 748 million people who do not have access to clean water, 325 million (43%) live in sub-Saharan Africa [11]. In Kenya, where 27% of the population obtain drinking water from an unimproved source [28], water scarcity is further exacerbated by the unequal geographic distribution of water resources [29,30] and large seasonal fluctuations in rainfall [31–33]. Within our study setting in Nyanza region, the mean rainfall during the wet season has been 180.05mm, with a range of 143 mm to 283.33 mm. A mean of 80.99 mm has been recorded for the dry season over a 10-year period (1990-2000), with a range of 18.24 to 110.46 mm [34].

In the course of our ongoing work on the consequences of food insecurity amongst pregnant and lactating women of mixed HIV status, formative research revealed that there were many experiences of water insecurity that were perceived to be deleterious [19,35]. For instance, formative work revealed that 77.3% of our participants felt ‘somewhat or strongly concerned’ for their physical safety during water acquisition; 64.1% reported drinking unsafe water; 65.3% reported worrying about accessing sufficient water and women without water in their compound spent an average of 4.5 (6.7) hours per week acquiring water [19]. Further, in qualitative work, women reported consequences of water insecurity including intimate partner violence, risk of miscarriage and stillbirth from carrying water, conflict with neighbors at water sources, and attacks from people and animals while fetching water [35].

A review of the extant literature showed a burgeoning field of household water insecurity scales developed in, e.g. Latin America [4,36], North America [37,38], South Asia [39], and sub-Saharan Africa [2,3,40]. However, these efforts to measure household water insecurity have been associated with a number of limitations including the lack of formative work, lack of robust scale validation, excessive length, and different target populations (i.e. not pregnant or post-partum) [1]. Also, the items that comprised the scale and analytic approaches used were highly variable across existing scales [1]. Furthermore, the dimensions of the existing scales were varied; while some scales showed a single dimension [2–4,36,40], others had multiple dimensions [39,41].

Further, we could find no scale for measuring household water insecurity validated for Kenya. Therefore, we set out to develop and validate a household water insecurity (HHWI) scale appropriate for pregnant and post-partum Kenyan women of mixed HIV status.

## Methodology

### Study setting and population

Data for scale development and validation were collected between June 2015 and August 2016 in Nyanza region, southwestern Kenya where the Luo, Kisi/Gusii, Kuria, and Luhya are the predominant ethnic groups. The major economic activities include fishing (on the nearby Lake Victoria), and mixed and agro-pastoral agriculture [42]. The region is typified by low crop yields and soil fertility, with a greater proportion of farmers engaged in subsistence farming [43]. Nyanza is one of the poorest regions in Kenya, with about 63% of the population living on less than $1 a day [44].

Research was conducted in the context of an observational cohort of 266 postpartum HIV-infected and HIV-uninfected women entitled “Pii en Ngima” [PEN], Luo for “water is life” (Clinicaltrials.gov# NCT02979418) who had been previously enrolled in a pregnancy cohort titled “Pith Moromo” (Clinicaltrials.gov # NCT02974972). HIV-infected women were over-sampled to achieve 1:1 ratio. The study took place in seven clinical catchment areas that span urban, peri-urban and rural sites across Nyanza region including Kisumu (urban), Migori (peri-urban), Nyahera (peri-urban), Rongo (peri-urban), Macalder (rural), Nyamaraga (rural), and Ongo (rural). Family AIDS Care and Education Services (FACES), an HIV care and treatment program in Nyanza region, supported each of the clinics in the medical sites.

Nyanza region was an appropriate study site because of the high level of food and water scarcity [19], the high prevalence of HIV, which is currently 6.9% for pregnant women in western Kenya [45], and the presence of an excellent clinical care, research and laboratory infrastructure through FACES.

### Data collection

Data collection for this study was structured in four phases (Table 1). The first phase, formative data collection, explored the experiences of water insecurity through “go-along interviews” (Activity A) [46,47], Photovoice photo elicitation interviews (Activity B) [48,49], and the Delphi method (Activity C) [50], which was conducted concurrently with focus group discussions (FGDs; Activity D)) [35]. The second phase involved the assembly (Activity E) and revision of HHWI scale questions using cognitive interviews (Activity F) [51–53]. The third phase entailed the administration of the survey to individual women (Activity G) and the final phase included collection of non-survey data for purposes of further scale validation (Activity H). Activities A, B, and D used non-cohort women (n=20) with similar demographic characteristics as those used in the third phase for survey administration, Activities G and H (n=241 and n=186, respectively).

**Table 1.** Data collection activities for the construction and development of the household water insecurity scale.

| Activity | Procedures | Purposes | Sample | Dates of activities |
| --- | --- | --- | --- | --- |
| **Phase 1: Formative data collection** |  |  |  |  |
| A. Go-along interviews of water access and use | Participant observation and HHWI<sup>a</sup> interview. | To explore experiences of household water use, acquisition and insecurity. | Non-cohort Kenyan women, n=20 | 06/2015-09/2015 |
| B. Photovoice (photo elicitation interviews) | Participants were briefly interviewed and lent digital cameras to take photographs of water related experiences. A second individual interview explored photographs and was followed by FGDs on most common emergent themes. | To explore experiences of household water use, acquisition, and insecurity. | Non-cohort Kenyan women, n=20 | 07/2015-10/2015 |
| C. The Delphi Method (S1 Fig., S1 Table)<sup>b</sup> | International experts on water and food insecurity were purposively selected to achieve a range of disciplines and geographic areas and asked to participate in online iterative surveys about HHWI | To identify and build consensus on key concepts related to HHWI | Non-cohort international professionals<br>Round 1, n=22<br>Round 2, n=17<br>Round 3, n=12 | 06/2015-01/2016 |
| D. Focus group discussions (FGDs) (S1 Fig) | After each Delphi round (Activity C), convenience sampling was used to select pregnant or postpartum field experts for FGDs in Kenya | To identify and build consensus on key concepts related to HHWI | Non-cohort Kenyan women<br>Round 1, n=15<br>Round 2, n=12 | 09/2015-11/2015 |
| **Phase 2: Assembly and Revision of HHWI Scale Questions** |  |  |  |  |
| E. Assembly of scale questions | Compiled initial HHWI questions based on steps A-D and existing literature. | To create an initial HHWI questionnaire | n=29 questions | 09/2015-11/2015 |
| F. Cognitive interviews | Questions from Activity E were asked, followed by probing questions | To determine if questions were understood as intended or could be improved. | Non-cohort Kenyan women, n=10 | 11/2015 |
| **Phase 3: Survey Administration at 15 & 18 months postpartum** |  |  |  |  |
| G. Household Water Insecurity survey module (S2 Table)<sup>c</sup> | Administered survey comprised of scale questions among mixed HIV status women | Measure HHWI in women's daily lives | PEN cohort participants<br>n=241 (15mpp<sup>d</sup>)<br>n=186 (18mpp) | 03/2016 – 09/2016<br>05/2016 – 10/2016 |
| H. Survey data for scale validation | Administered survey questions about time spent collecting water, the primary source of drinking water, amount of money spent purchasing water, individual food insecurity, and perceived stress | To validate HHWI scale | PEN cohort participants<br>n=241 (15mpp)<br>n=186 (18mpp) | 03/2016 – 09/2016<br>05/2016 – 10/2016 |
| **Phase 4: Non-survey data for further validation** |  |  |  |  |
| I. Drinking water quality | Measured *Escherichia coli* concentrations using Colilert™ and Compartment Bag Test (CBT) assays | Measure water quality | PEN cohort women, n=35 | 01/2016 |
| J. Water quantity (stored water and amount of water used) | Measured the quantity of drinking water stored and used by the household (in liters) | Measure total household drinking water stores and total household water use | PEN cohort women, n=35 | 01/2016 |
| K. Retrospective Recall | Two exercises were conducted with a randomly selected subset of respondents. The first was administered daily for 30 days and the second administered retrospectively on the 31<sup>st</sup> day. | Data collected to assess intra-respondent reliability | PEN cohort women, n=35 | 11/2016 |

Notes: <sup>a</sup>household water insecurity; <sup>b</sup>Supplementary Table 1; <sup>c</sup>Supplementary Table 2; <sup>d</sup>months postpartum

### Phase 1: Formative data collection

Although the results of our Phase 1 are presented elsewhere to avoid an excessively long manuscript [35], we briefly describe the formative methods used in order to convey the basis of the initial scale questions and to place our scale development activities within a broader context.

#### A. Go-along interviews

Go-along interviews are a hybrid between participatory research and qualitative in-depth interviewing, that attempt to contextualize meaning within social and spatial contexts [46,47,54]. In this study, a Kenyan anthropologist (PM) accompanied participants to and from water collection sites while asking questions. The interviews were translated (from Swahili or Luo to English), transcribed, coded and analyzed using Dedoose software (Los Angeles, CA: SocioCultural Research Consultants, LLC).

#### B. Photovoice

Photovoice applies documentary photography and critical dialogue to explore the lived experiences of people and as a means of sharing knowledge [48,49]. In this study, twenty women were lent digital cameras to take photos of their experiences of household water acquisition, use and insecurity. On a second visit, these photos were used to conduct in-depth individual interviews. A subset of these photos became the core focus for dialogues about HHWI during FGDs at a third encounter. Dedoose was also used to code translated transcripts from in-depth interviews and FGDs.

#### C. Delphi method

The Delphi method is a technique “for structuring a group communication process so that the process is effective in allowing a group of individuals, as a whole, to deal with a complex problem” [50]. Here, it was used to obtain feedback from international experts including those with expertise in hydrology and geographic research, WASH and water related programs, policy implementation, food insecurity and scale development, over the course of three rounds of surveys (**S1 Fig**). Each round was interspersed with FGDs in which questionnaires progressively became more closed ended. Questions included the definition of water insecurity, household water-related activities, barriers to water acquisition, consequences of water insecurity, and possible survey items that could constitute a HHWI scale (**S1 Table**).

#### D. Focus group discussions

FGDs were conducted iteratively with the Delphi process (S1 Fig.). To participate in FGDs, nurses and healthcare professionals purposively recruited postpartum women who were available and were either pregnant or had children less than 2 years of age in 4 study areas. After Delphi round 1, FGD participants (Kisumu; n=8 and Rongo; n=7) were asked to provide feedback on topics discussed in the online survey to build consensus around the definition of and questions related to HHWI. Another group of FGD participants (Migori; n=5 and Macalder; n=7) also provided information with which to revise questions for the survey.

### Assembly and Revision of Scale Questions

#### E. Assembly of scale questions

Based on themes that emerged from activities A-D [35], and the burgeoning literature on measurement of household water insecurity [2–4,37,36], we created 29 questions related to HHWI. Thirteen (13) of the questions (marked with asterisk in S2 Table) on the psychological, social, economic, and health consequences of water insecurity were adapted and modified from previous water insecurity scales [2,3,36,40]; the other 16 questions originated from activities A-D. The 29 questions were then ordered in what we considered to be least to most severe manifestations of water insecurity. They were phrased as Likert-type items, with the following response options: never (0), rarely (1-2 times), sometimes (3-10 times), and often (more than 10 times) in the last four weeks.

#### F. Cognitive interviewing

Once the questions were developed, we conducted cognitive interviews (n=10). Cognitive interviewing was used to assess: whether participants perceived the intent of the water insecurity questions as intended, whether participants were able to repeat questions they had been asked and the thought processes behind their responses, and whether the response options were appropriate and/or adequate [51–53]. This interviewing approach resulted in only some minor rephrasing of the 29 items (**S2 Table**).

### HHWI Survey Administration (15 & 18 months postpartum)

#### G. HHWI scale questions

The HHWI module was then administered as part of the PEN study at 15 and 18 months postpartum.

#### H. Survey data for scale validation

Participants were also asked other survey questions pertaining to water and their physical and psychological health for purposes of scale validation.

Water acquisition questions were asked to help assess convergent validity. Participants were asked to indicate how long it took for them to travel to the water source, queue, fetch water and return to their houses and how much they spent on water. Additionally, we assessed access to safe water by using the WHO/UNICEF [11] survey questions for improved and unimproved drinking water sources.

Food insecurity was assessed for purposes of predictive validity. We used the Individual Food Insecurity Access Scale (IFIAS) [51], which is a 9-item scale analogous to the Household Food Insecurity Access Scale (HFIAS) [55] but asks participants about their own individual experiences with access to food in the prior month. The intensity of food insecurity was assessed with follow-up questions asking whether this condition was experienced never, rarely, sometimes, or often (coded 0, 1, 2 or 3) with a range of 0-27.

Maternal stress was also assessed to examine predictive validity. We used Cohen’s Perceived Stress Scale [56], These questions asked about the feelings and thoughts of women in the prior month, i.e. the frequency they felt upset, nervous or worried. The intensity of perceived stress was assessed by Likert-type response format of never, almost never, sometimes, fairly often and very often (coded 0, 1, 2, 3 or 4) with a range of 0-40.

### Non-survey data for further validation

#### I. Water quality

Data on water quality were collected at 5 randomly selected PEN participant households in each of the 7 catchment areas (n=35). Drinking water quality was assessed by aseptically collecting triplicate 100ml water samples using Whirl-Pak Thio-Bags to test for coliform and *E.coli* MPN using Compartment Bag Tests (CBT) (Aquagenx) and Colilert (Idexx Laboratories) [57]. Samples collected were analyzed for total coliform most probable number (MPN) and *E.coli* MPN using CBT and Colilert methodologies. Water quality was dichotomized according to WHO standards showing the presence of *Escherichia coli* (≥1 MPN/100 ml) in household drinking water [58–60].

#### J. Water quantity

In the same 35 households, we measured the amounts of stored water for drinking and non-drinking purposes at a single time point in a given day. The volume was measured in litres based on the size of the storage containers and the amount of water in the container. For instance, a half-full 20-litre jerry can was measured as 10 litres of stored water. We also measured the amount of water used daily by the household in litres based on estimates of the amount of water used in cooking, drinking, washing foods, washing clothes, bathing, washing face, brushing teeth, washing hands, washing utensils/dishes, and washing toilets. By dividing the total amount of water used by number of individuals in the household, we were able to estimate per capita household daily water use. For purposes of analysis, complete data were available for 27 households. Of the eight households dropped, 3 had no stored drinking water and 5 had data available from a different time point.

#### K. Retrospective recall

To assess intra-respondent reliability, we administered a subset of the 29 items (20-item version of the water insecurity module) daily for 30 days. We used 20 items to reduce respondents’ fatigue as it was being asked continuously for a month. Thirty-five participants were asked each day if they had that experience of water insecurity in the prior day and could respond yes or no. On the 31^st^ day, participants were asked to indicate the number of days they had experienced that particular aspect of HHWI over the prior 30 days. Correlation coefficients were calculated between cumulative daily recall and responses from the 31^st^ day.

## Data Analyses

Quantitative data analyses were conducted in six phases including descriptive analyses, item reduction, extraction of factors, tests of dimensionality, scale reliability, and validity (Table 2). Software packages used included M*Plus* version 7.40 (Los Angeles, CA: Muthén & Muthén) [61], SPSS version 20.0 (Armonk, NY: IBM Corp.) and STATA version 14 (College Station, TX: StataCorp LP). Tests of dimensionality were conducted using data from 18 months postpartum (n=186); the rest of the analyses were done using data from 15 months postpartum (n=241).

**Table 2.** Analytical procedures for the construction and development of household water insecurity scale among postpartum women in western Kenya.

| Concept | Purpose | How assessed | References |
| --- | --- | --- | --- |
| **A. Descriptive Analyses** |  |  |  |
| Summary statistics | To examine the distribution of all scale items and measured variables. | Conducted a summary statistics of scale items and all measured variables relevant to HHWI. |  |
| **B. Item Reduction** |  |  |  |
| Adequate Variance | To examine the variation between items in HHWI scale | Analyzed the distribution of items for HHWI. | [62] |
| Polychoric Correlations (Inter-item) | To determine the correlations between scale items. | Estimated average inter-item correlation coefficient, help to determine which items to drop. | [61,63,64] |
| Polyserial Correlations (Item-total) | To determine the correlation between individual scale items with the sum score of all scale items. | Estimated adjusted item-total correlation coefficients, help to determine which items to drop. | [61,63,64] |
| Item Communalities | To determine the measurement error in each item or the true score variance. | Estimated using principal axis factoring. | [62] |
| Kaiser-Meyer-Olkin (KMO) Test for sampling adequacy | To measure the proportion of common variance among items and determine whether the data is suitable for factor analysis. | Estimated the sampling adequacy for each item in the model and for the complete model. KMO values between 0.8 and 1 indicate the sample is adequate. | [65] |
| Bartlett Test of Sphericity | To compare the observed correlation matrix to the identity matrix. | Tested the null hypothesis that the correlation matrix has an identity matrix. | [66–68] |
| **C. Extraction of Factors** |  |  |  |
| Exploratory Factor Analysis (latent Structure) | To measure the structure of a set of observed variables and identify the subset of variables that corresponds to each of the underlying dimensions. | Factor analysis of retained twenty items used together with the Guttman-Kaiser >1 rule and Cattell's scree plot. | [61,64,69–72] |
| Parallel Analysis | To identify the possible number of factors that can be developed from the data. | Estimated number of identifiable factors from scale items. This was a form of sensitivity analysis to the exploratory factor analysis. | [73,61,64,70] |
| Model Fit Assessment | To determine the fitness of both factor and parallel analyses to the data. | Examined model fit indices against acceptable thresholds (**S3 Table**). | [74–81] |
| **D. Tests of Dimensionality** |  |  |  |
| Confirmatory Factor Analysis (Latent Variable Modeling) | To address queries on the latent structure of scale items and their underlying relationships. i.e. to validate previous EFA results. | Factor analysis of items via Structural Equation Modeling. This also helped with the determination of construct validity of the HHWI scale. | [61,69,82,83] |
| Bifactor Analysis | To evaluate dimensionality-related questions. | With bifactor analysis, the factor loadings of the general factor were compared to the group factors to help determine the dimensionality of the scale. | [84–86] |
| Model Fit Assessment | To determine the fitness of both confirmatory factor analysis and bifactor modeling solutions. | Examined model fit indices against acceptable thresholds (**S3 Table**) | [74–81] |
| **E. Scale Reliability** |  |  |  |
| Intra-respondent reliability | To assess the stability and consistency of responses by respondents on scale items. | Correlated the sum score of daily retrospective responses on HHWI items for 30 days with scores on a 30-day recall. | [87] |
| Coefficient alpha | To assess the internal consistency of the scale. i.e., the degree to which the set of items in the scale co-vary, relative to their sum score. | Calculated Cronbach's alpha for scale items at 15 months postpartum and 18 months postpartum. | [88] |
| Coefficient of stability | To assess the degree to which the participant's performance is repeatable; i.e. how consistent their scores are across time. | Estimated the coefficient of stability via Test-retest reliability. This was indexed by the correlation coefficient of two assessments of HHWI at two different time points. | [63,64,89] |
| **F. Scale Validity** |  |  |  |
| Predictive validity | To determine the degree to which test scores predict criterion measurements to be made in the future. | Estimated the association between HHWI and maternal stress to food insecurity scores. | [63,64] |
| Convergent validity | To examine the evidence that the same concept measured in different ways yields similar results. | Estimated the correlation between HHWI and water quality (*E.coli* concentrations), time to water collection, amount spent on water for household, and season of data collection. | [89–91] |
| Discriminant validity | To examine the evidence that the concept measured is different from other closely related concepts. | Estimated the correlation between HHWI and per capita household water use. Indicated by predictably low correlations between HHWI and other measures. | [89–91] |
| Differentiation by 'known groups' | To examine the degree to which the concept measured behaves as expected in relation to 'known groups'. | Estimated a differential test of means for maternal HIV status, season, water quality, and source of drinking water. | [89–91] |

### A. Descriptive Analyses

First, we estimated proportions, means, and standard deviations of the HHWI module and participant characteristics. Although there were 5-response categories for the scale items originally, the sample distribution was skewed to the right (<5%) for “always” for each item. Therefore, “often” and “always” were collapsed for subsequent analyses.

### B. Item Reduction

We first assessed adequate variance for all HHWI items [62]. This was followed by polychoric (inter-item) and polyserial (item-total) correlation of scale items [61,63,64]. Items without adequate variance, very low inter-item (<0.3) and item-total correlations (<0.3), very high residual variances (>0.50), and high missing cases (>10%) were dropped. We also estimated item communalities for degree of common variance between items [62], the Kaiser-Meyer-Olkin measure for sampling adequacy [65], and the Bartlett test of sphericity [66–68] to ensure our item reduction approach was robust. Furthermore, one item was dropped for any two items that suggested collinearity (≥0.98).

### C. Extraction of Factors

Multiple approaches were used to determine the number of factors to retain. Exploratory factor analysis (EFA) was used together with Guttman’s [92] eigenvalue rule of lower bound, Kaiser’s [71] eigenvalue >1 rule, Cattell’s [72] scree test, and Horn’s [73] parallel analysis (PA) to determine the optimal number of factors that fit the data at 15 months postpartum [73,61,64,70]. For the scree tests, the root of the scree was used as a point of extraction for the true number of factors [72]. The extraction process in all the models used oblique rotation with weighted least squares with mean and variance adjustment (WLSMV) estimator except for Horn’s PA, which employed the maximum likelihood (ML) estimator. For sensitivity analysis, we employed principal axis factors.

A number of model fit statistics were used to determine meaningful model fitness for both traditional factor and parallel analyses (**S3 Table**). The fit indices included Chi-square test of model fit, the Tucker Lewis Index (TLI≥0.95), the Comparative Fit Index (CFI≥0.95), the Root Mean Square of Error of Approximation (RMSEA≤0.10), and the Standardized Root Mean Square Residual (SRMR≤0.08) [74–81]. Consistent with the factor structure of previous household water insecurity scales elsewhere [2–4,36,40], we assumed our model will produce similar factor structure for our scale.

### D. Tests of Dimensionality

In order to test the factor structure obtained from the EFA, a test of scale dimensionality was conducted using confirmatory factor analysis (CFA) and bifactor or nested factor modeling on an independent sample at 18 months postpartum. The CFA allows for the test of dimensionality of the hypothesized factors [61,69,82,83]. Complementarily, the bifactor model allows researchers to extract a primary unidimensional construct while recognizing the multidimensionality of the construct [84–86]. The bifactor model assumes each item loads on two dimensions. The first is a general latent trait or factor that underlies all the scale items and the second, a group factor. This approach allows researchers to examine any distortion that may occur when unidimensional CFA models are fit to multidimensional data [84–86].

To determine whether to retain a construct as unidimensional or multidimensional, the factor loadings from the general factor are compared to those from the group factors (sub scales) [85,86]. Where the factor loadings on the general factor are significantly larger than the group factors, a unidimensional scale is implied [85,93]. The model fitness of both the confirmatory factor and bifactor models were assessed using TLI, CFI, RMSEA and the Weighted Root Mean Square Residual [74–81] (**S3 Table**).

### E. Scale Reliability

We assessed intra-respondent reliability of scale items retrospectively by comparing daily recall across 30 days with the sum score of a retrospective recall on the 31^st^ day. This was to assess the stability and consistency of responses on scale items.

The reliability of the scale itself was estimated using coefficient alpha and the coefficient of stability. First, Cronbach’s coefficient alpha of scale items was calculated for the samples at 15 and 18 months postpartum to compare and correlate the observed score variation between each of the items in the scales for both samples [81]. Second, we assessed the coefficient of stability (test-retest reliability), which involved the correlation of scale scores at 15 and 18 months postpartum [63,64,89].

### F. Scale Validity

We used predictive (criterion), construct (convergent and discriminant) validity and differentiation between ‘known groups’ to assess scale validity. Predictive (criterion) validity was assessed by examining the associations between HHWI and perceived maternal stress as well as food insecurity [56,57].

Convergent validity was measured against time to and from water source and amount of money spent on purchasing water in the past month. We calculated Pearson product-moment correlations based on Fisher’s transformation [89–91].

Discriminant validity was assessed by correlating HHWI with per capita water used daily [89–91], which has similarly been used in previous studies [2,3,36]. Consistent with the findings of Tsai et al [3], Hadley and Wutich [36], and Stevenson et al. [2], we assumed that there would be little or no relationship between HHWI and per capita water used, i.e. that HHWI is distinct from household water *use*.

As a final measure of validity, we assessed the scale score by differentiating the position of ‘known groups’. In other words, we expected to have significantly higher HHWI scores for participants whose water was contaminated with *Escherichia coli (E.coli)*, were HIV positive, those who used unimproved sources of water, and during the dry season. We used Student’s *t*-test for this analysis.

### IRB Approval

We obtained approval for this study from the Institutional Review Boards at Cornell University, Northwestern University, and the Kenyan Medical Research Institute Scientific and Ethics Review Committee. Also, we obtained written informed consent from all participants in this study.

## Results

### Initial household water insecurity module

Formative work in Phase 1 resulted in the creation of 29 HHWI questions (Activity E, Table 1). The cognitive interviews (Activity F) indicated people were able to understand the intended meaning and could accurately repeat the questions. Responses from cognitive interviews resulted in the adjustment of the structure of the questions and the retraining of interviewers on how to ask questions without ambiguity and the avoidance of leading prompts. The response options that were considered appropriate for a 4-week recall period were “never” (0), “rarely” (meaning 1-2 times), “sometimes” (3-10 times), “often” (10-20 times) and “always” (>20 times).

### Participant Characteristics

Of the 241 participants who were interviewed at 15 months postpartum, the mean household size was 3.5 with a range of 1-12 members (Table 3). The majority of women (90.5%) interviewed were primiparous, 51.5% were HIV positive, and the mean age was 25 (range 18-39) years. Participants who lived in rural (35.3%) and peri-urban (21.6%) regions comprised the majority of the sample. Consistent with the general demography of Nyanza region, 52.5% of the sample had primary education and 8.8% had college education. The mean Individual Food Insecurity Access Score was 5.4 (SD=5.5) with a Cronbach’s alpha of 0.90. Participants had a mean perceived stress score of 17.3 (SD=4.3) [94], with a Cronbach’s alpha of 0.70. For the 15-month postpartum visit, most of the participants (62.2%) were interviewed during the rainy season (Table 3).

**Table 3.**
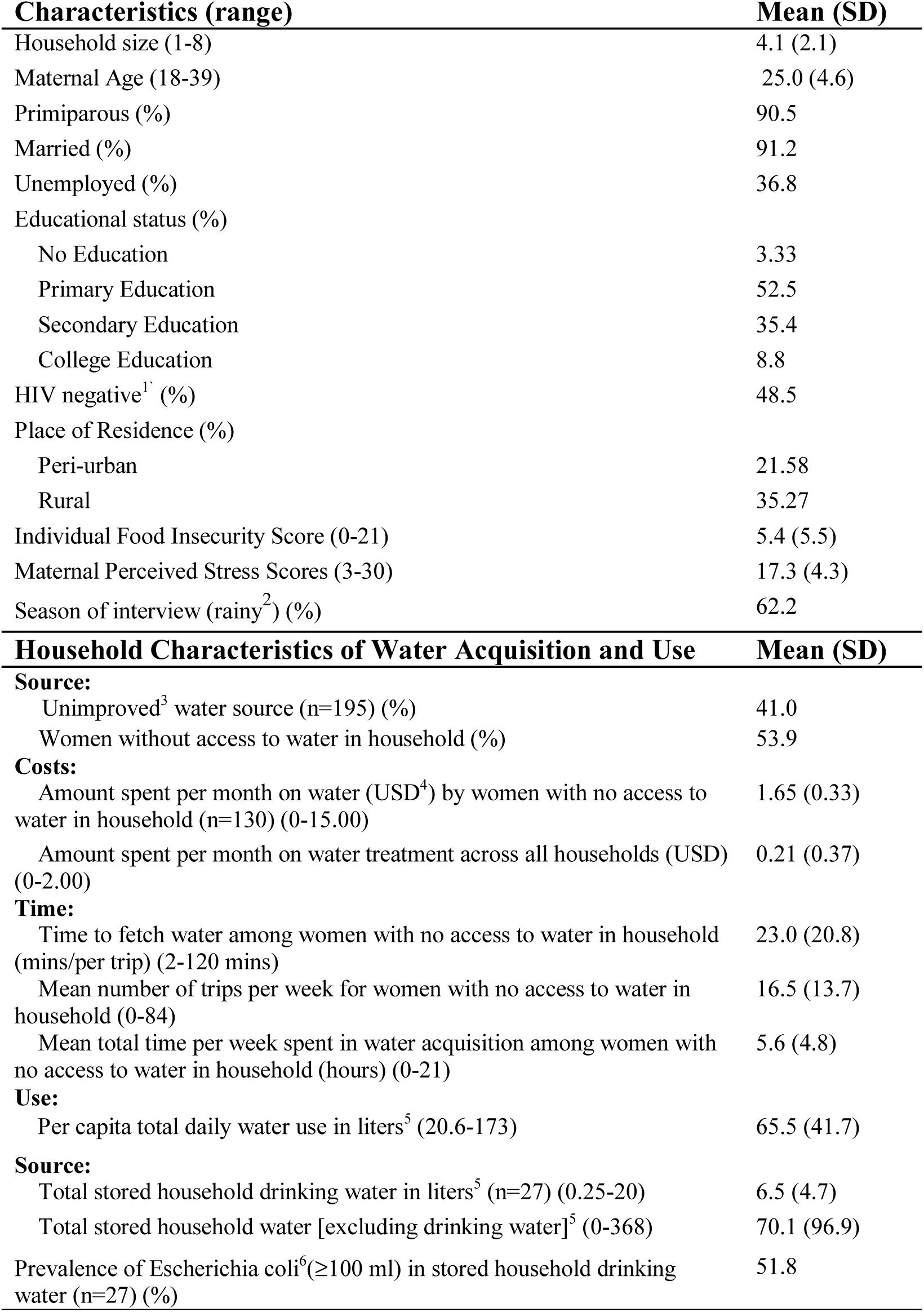
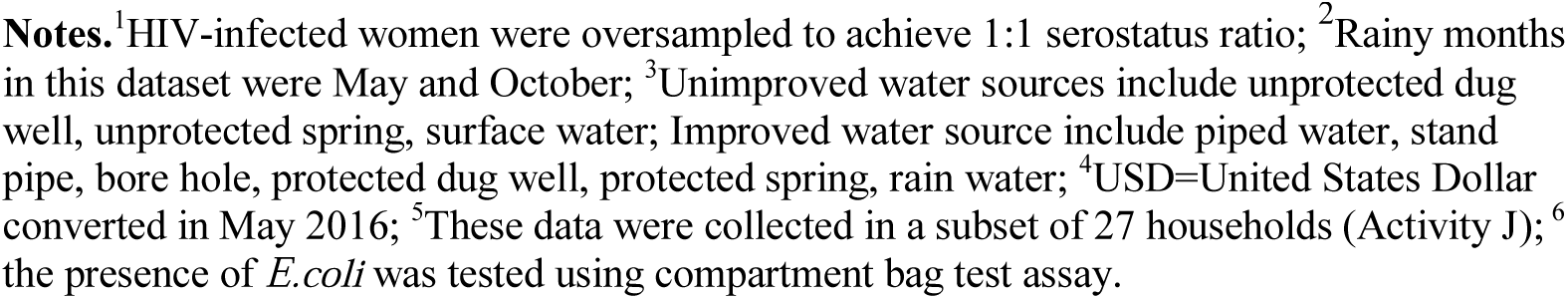
Socio-demographic, water access and use among Kenyan Women of mixed HIV status at 15 months postpartum (N=241)

| **Characteristics (range)** | **Mean (SD)** |
| --- | --- |
| Household size (1-8) | 4.1 (2.1) |
| Maternal Age (18-39) | 25.0 (4.6) |
| Primiparous (%) | 90.5 |
| Married (%) | 91.2 |
| Unemployed (%) | 36.8 |
| Educational status (%) |  |
| No Education | 3.33 |
| Primary Education | 52.5 |
| Secondary Education | 35.4 |
| College Education | 8.8 |
| HIV negative<sup>1</sup> (%) | 48.5 |
| Place of Residence (%) |  |
| Peri-urban | 21.58 |
| Rural | 35.27 |
| Individual Food Insecurity Score (0-21) | 5.4 (5.5) |
| Maternal Perceived Stress Scores (3-30) | 17.3 (4.3) |
| Season of interview (rainy<sup>2</sup>) (%) | 62.2 |
| **Household Characteristics of Water Acquisition and Use** | **Mean (SD)** |
| **Source:** |  |
| Unimproved<sup>3</sup> water source (n=195) (%) | 41.0 |
| Women without access to water in household (%) | 53.9 |
| **Costs:** |  |
| Amount spent per month on water (USD<sup>4</sup>) by women with no access to water in household (n=130) (0-15.00) | 1.65 (0.33) |
| Amount spent per month on water treatment across all households (USD) (0-2.00) | 0.21 (0.37) |
| **Time:** |  |
| Time to fetch water among women with no access to water in household (mins/per trip) (2-120 mins) | 23.0 (20.8) |
| Mean number of trips per week for women with no access to water in household (0-84) | 16.5 (13.7) |
| Mean total time per week spent in water acquisition among women with no access to water in household (hours) (0-21) | 5.6 (4.8) |
| **Use:** |  |
| Per capita total daily water use in liters<sup>5</sup> (20.6-173) | 65.5 (41.7) |
| **Source:** |  |
| Total stored household drinking water in liters<sup>5</sup> (n=27) (0.25-20) | 6.5 (4.7) |
| Total stored household water [excluding drinking water]<sup>5</sup> (0-368) | 70.1 (96.9) |
| Prevalence of Escherichia coli<sup>6</sup>(≥100 ml) in stored household drinking water (n=27) (%) | 51.8 |

**Notes.**<sup>1</sup>HIV-infected women were oversampled to achieve 1:1 serostatus ratio; <sup>2</sup>Rainy months in this dataset were May and October; <sup>3</sup>Unimproved water sources include unprotected dug well, unprotected spring, surface water; Improved water source include piped water, stand pipe, bore hole, protected dug well, protected spring, rain water; <sup>4</sup>USD=United States Dollar converted in May 2016; <sup>5</sup>These data were collected in a subset of 27 households (Activity J); <sup>6</sup> the presence of *E.coli* was tested using compartment bag test assay.

### Water access and use

Of the 241 participants interviewed, nearly half (41.0%) used drinking water from unimproved sources, and 53.9% did not have access to water in their households or compounds. Of the women who had access to water in their households, 8.8% were unimproved sources. Women with no access to water in their households spent a mean amount of 1.60 USD; with a range of 0 to 15.00 US dollars a month on water acquisition. The mean amount spent on water treatment across all households was 0.21 USD. Women without access to water in their households spent a mean time of 23.0 minutes per trip and 16.5 trips per week acquiring water for their households, for a mean of 5.6±4.8 hours per week. In 27 households in which we had data to assess water use and microbial analysis, a mean of 65.5 liters of water was used daily by households, 6.5 liters were stored for drinking, and a mean of 70.1 were stored for other uses. For microbial analysis, 51.8% (14 out of 27) of the households tested positive (≥100 ml) for E*.coli* in stored drinking water (Table 3).

The most severe manifestations of water insecurity, such as sleeping thirsty and having no water in the household whatsoever, were least endorsed (23. 8% and 19.2%) in this population (Fig. 1, Table 4). Items that reflected less severe expressions of water insecurity, including worrying about having enough water and drinking water that was considered to be unsafe, were considerably more common (46.3% and 44.9% respectively), with 3.7% experiencing these events often or always (Fig.1, Table 4).

**Fig 1.**
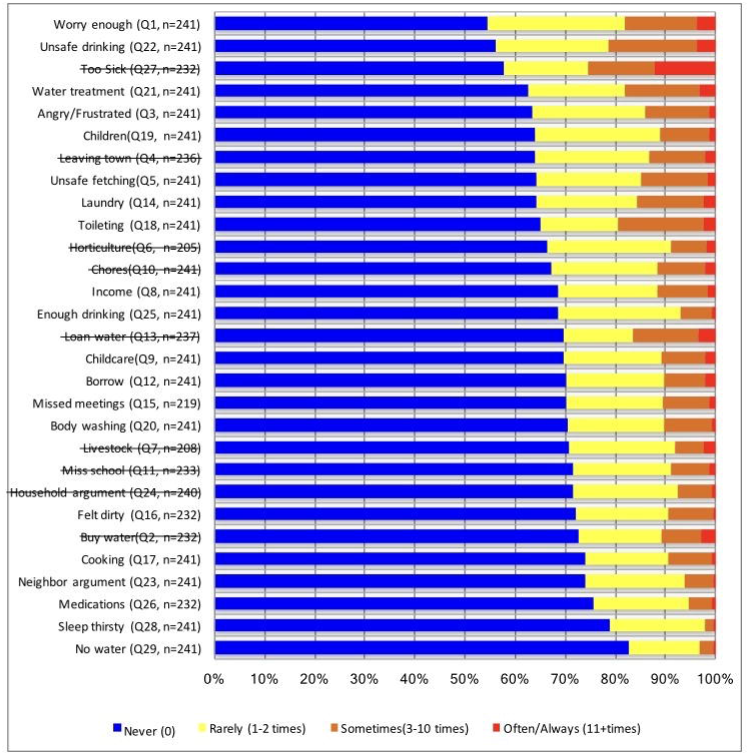
The distribution of response categories for all 29 Household Water Insecurity Scale Items at 15 months postpartum with total responses (n=241).

**Table 4.**
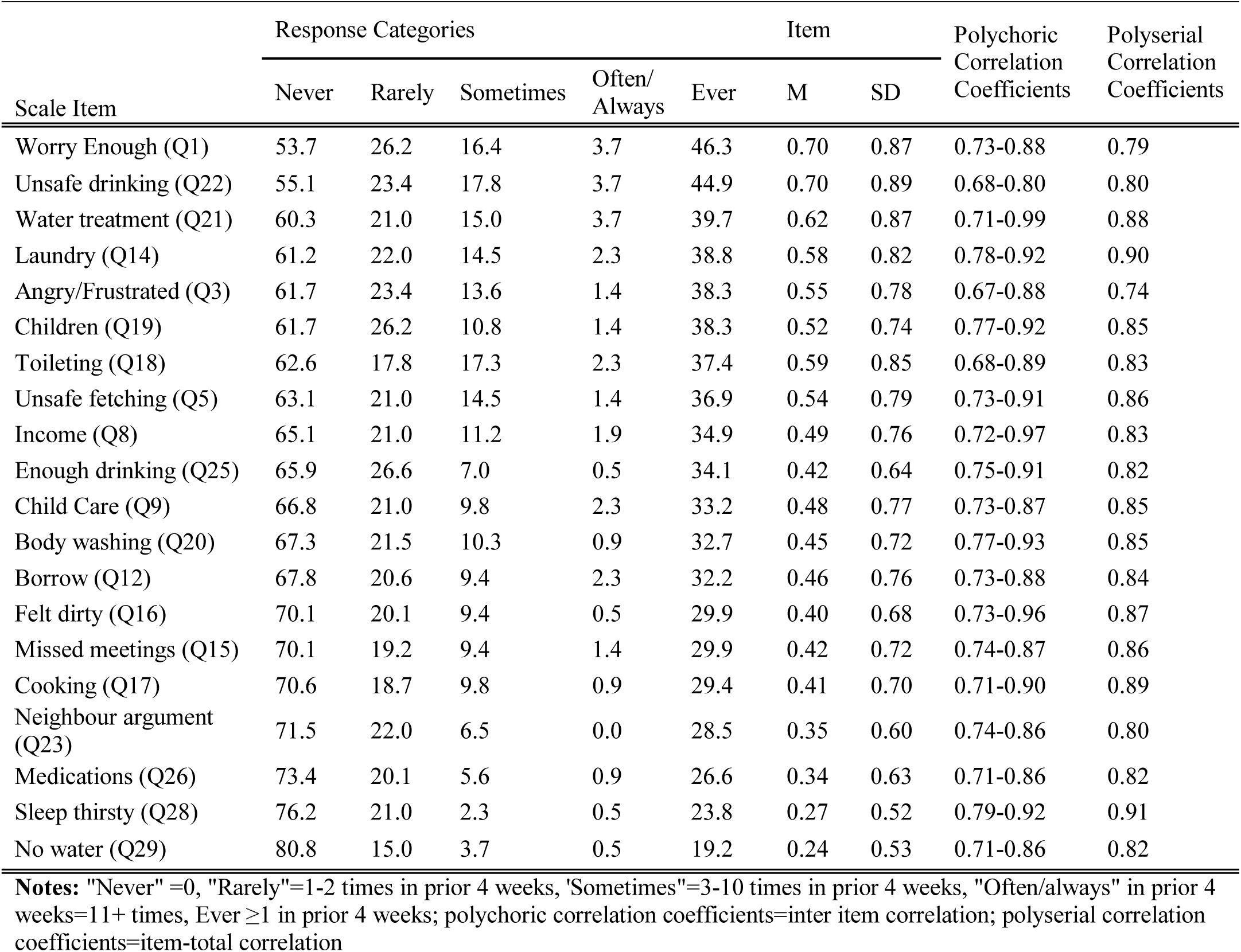
Frequency distribution of response categories and polychoric/polyserial correlation coefficients for household water insecurity items among women in western Kenya at 15 months postpartum, from highest to the lowest frequency (n=241).

| Scale Item | Response Categories |  |  |  |  | Item |  | Polychoric<br>Correlation<br>Coefficients | Polyserial<br>Correlation<br>Coefficients |
| --- | --- | --- | --- | --- | --- | --- | --- | --- | --- |
|  | Never | Rarely | Sometimes | Often/<br>Always | Ever | M | SD |  |  |
| Worry Enough (Q1) | 53.7 | 26.2 | 16.4 | 3.7 | 46.3 | 0.70 | 0.87 | 0.73-0.88 | 0.79 |
| Unsafe drinking (Q22) | 55.1 | 23.4 | 17.8 | 3.7 | 44.9 | 0.70 | 0.89 | 0.68-0.80 | 0.80 |
| Water treatment (Q21) | 60.3 | 21.0 | 15.0 | 3.7 | 39.7 | 0.62 | 0.87 | 0.71-0.99 | 0.88 |
| Laundry (Q14) | 61.2 | 22.0 | 14.5 | 2.3 | 38.8 | 0.58 | 0.82 | 0.78-0.92 | 0.90 |
| Angry/Frustrated (Q3) | 61.7 | 23.4 | 13.6 | 1.4 | 38.3 | 0.55 | 0.78 | 0.67-0.88 | 0.74 |
| Children (Q19) | 61.7 | 26.2 | 10.8 | 1.4 | 38.3 | 0.52 | 0.74 | 0.77-0.92 | 0.85 |
| Toileting (Q18) | 62.6 | 17.8 | 17.3 | 2.3 | 37.4 | 0.59 | 0.85 | 0.68-0.89 | 0.83 |
| Unsafe fetching (Q5) | 63.1 | 21.0 | 14.5 | 1.4 | 36.9 | 0.54 | 0.79 | 0.73-0.91 | 0.86 |
| Income (Q8) | 65.1 | 21.0 | 11.2 | 1.9 | 34.9 | 0.49 | 0.76 | 0.72-0.97 | 0.83 |
| Enough drinking (Q25) | 65.9 | 26.6 | 7.0 | 0.5 | 34.1 | 0.42 | 0.64 | 0.75-0.91 | 0.82 |
| Child Care (Q9) | 66.8 | 21.0 | 9.8 | 2.3 | 33.2 | 0.48 | 0.77 | 0.73-0.87 | 0.85 |
| Body washing (Q20) | 67.3 | 21.5 | 10.3 | 0.9 | 32.7 | 0.45 | 0.72 | 0.77-0.93 | 0.85 |
| Borrow (Q12) | 67.8 | 20.6 | 9.4 | 2.3 | 32.2 | 0.46 | 0.76 | 0.73-0.88 | 0.84 |
| Felt dirty (Q16) | 70.1 | 20.1 | 9.4 | 0.5 | 29.9 | 0.40 | 0.68 | 0.73-0.96 | 0.87 |
| Missed meetings (Q15) | 70.1 | 19.2 | 9.4 | 1.4 | 29.9 | 0.42 | 0.72 | 0.74-0.87 | 0.86 |
| Cooking (Q17) | 70.6 | 18.7 | 9.8 | 0.9 | 29.4 | 0.41 | 0.70 | 0.71-0.90 | 0.89 |
| Neighbour argument (Q23) | 71.5 | 22.0 | 6.5 | 0.0 | 28.5 | 0.35 | 0.60 | 0.74-0.86 | 0.80 |
| Medications (Q26) | 73.4 | 20.1 | 5.6 | 0.9 | 26.6 | 0.34 | 0.63 | 0.71-0.86 | 0.82 |
| Sleep thirsty (Q28) | 76.2 | 21.0 | 2.3 | 0.5 | 23.8 | 0.27 | 0.52 | 0.79-0.92 | 0.91 |
| No water (Q29) | 80.8 | 15.0 | 3.7 | 0.5 | 19.2 | 0.24 | 0.53 | 0.71-0.86 | 0.82 |

**Notes:** "Never" =0, "Rarely"=1-2 times in prior 4 weeks, 'Sometimes'=3-10 times in prior 4 weeks, "Often/always" in prior 4 weeks=11+ times, Ever $\geq 1$ in prior 4 weeks; polychoric correlation coefficients=inter item correlation; polyserial correlation coefficients=item-total correlation

### Item Reduction

In total, nine scale items were dropped from the 29-question survey (Fig. 1). We first dropped two questions related to horticulture and livestock because they had negative and weak inter-item (<0.3) and item total (<0.3) correlation coefficients. Also, questions on both horticulture (13.7%) and livestock (14.9%) had high missing cases. Further, six items (Leaving Town, Miss School, Household Arguments, Loan Water, Too Sick, and Buy Water) were dropped due to their very large residual variances (>0.5) and very low communalities (<0.3). A final item, Chores, was dropped because its very high correlation (*r*=0.98) with ‘Childcare’ created redundancy in items.

We then investigated inter-item (polychoric) and item-total (polyserial) correlations (Table 4). Inter-item correlations were strong, ranging from 0.67 to 0.97 for the remaining 20 items. Polyserial coefficients also showed very strong item-total correlations, ranging from 0.74 to 0.90, with an average item-total correlation of 0.84. Our Sensitivity tests showed the communalities in the 20 remaining items were all above 0.69, suggesting each of the items had some common variance with other items. The Kaiser-Meyer-Olkin measure of sampling adequacy was 0.94, above the recommended value of 0.60, and the Bartlett’s sphericity test was significant (χ^2^ (190) = 4436.25, p < .001). These indicators suggested that all 20 items should be used to explore the number of factors behind the correlation matrix [64,67,68].

### Extraction of Factors

To understand the latent factor structure of our items, we used EFA and the Guttman-Kaiser rule to extract two factors from the data with initial eigenvalues of 15.86 for factor one and 1.02 for factor two (Table 5). This was confirmed by Horn’s parallel analysis with eigenvalues 13.33 and 2.24 for factors one and two respectively. However, the amounts of variation explained by the first factor in both analyses were much bigger (79.3% and 66.7% respectively) suggesting a single underlying factor for HHWI [85]. Further, an evaluation of scree plots in both analyses showed a single dominant factor (**S2 Fig**). Specifically, the line for the average simulated eigenvalue was above the empirical factor 2 eigenvalue **(S3 Fig)**. This also suggested a one-factor solution was most appropriate.

**Table 5.** Model fit indices of factor extraction at 15 months postpartum and tests of dimensionality at 18 months postpartum.

| Factor Extraction (N=241) |  |  |  |  |  |  |  |
| --- | --- | --- | --- | --- | --- | --- | --- |
| Rotation | Analytical Technique | $\chi^2$<sup>3</sup> | df<sup>4</sup> | RMSEA<sup>5</sup> | CFI<sup>6</sup> | TLI<sup>7</sup> | SRMR<sup>8</sup> |
| Geomin Oblique | EFA<sup>1</sup>: HHWI (20 items) |  |  |  |  |  |  |
|  | 1 Factor (Ev<sup>2</sup>=15.86) | 1144 | 170 | 0.15 | 0.97 | 0.97 | 0.06 |
|  | 2 Factors (Ev=1.02) | 526.1 | 151 | 0.10 | 0.99 | 0.99 | 0.04 |
| Tests of Dimensionality (N=186) | | $\chi^2$ | df | RMSEA | CFI | TLI | WRMR<sup>9</sup> |
| Geomin Oblique | CFA<sup>10</sup> (1 Factor) | 1079.4 | 170 | 0.17 | 0.96 | 0.96 | 2.05 |
| Bi-Geomin Oblique | Bifactor | 412.9 | 150 | 0.09 | 0.99 | 0.98 | 0.94 |

**Notes:** <sup>1</sup>Exploratory Factor Analysis; <sup>2</sup>Eigenvalues; <sup>3</sup>chi-square goodness of fit statistic; <sup>4</sup>degrees of freedom; <sup>5</sup>RMSEA ( $\leq 0.10$ ) = Root Mean Square Error of Approximation; <sup>6</sup>CFI ( $> 0.95$ )=Comparative Fit Index; <sup>7</sup>TLI ( $> 0.95$ )=Tucker Lewis Index; <sup>8</sup>SRMR ( $\leq 0.08$ )=Standardized Root Mean Square Residual; <sup>9</sup>WRMR ( $< 1.0$ )=Weighted Root Mean Square Residual; <sup>10</sup>Confirmatory Factor Analysis. All chi-square goodness-of-fit tests were statistically significant at $p < 0.001$ .

An evaluation of the factor loadings associated with the eigenvalues produced two solutions, a one-factor model and two-factor model (Table 6). An examination of the factor loadings for the two-factor model showed three statistically significant cross loading items [toileting (0.75 vs. 0.32), unsafe drinking (0.52 vs. 0.68), and water treatment (0.70 vs. 0.50)]. However, the scores on the dominant factor were comparatively higher than the second factor, supporting a single dominant factor [84,85,95].

**Table 6.**
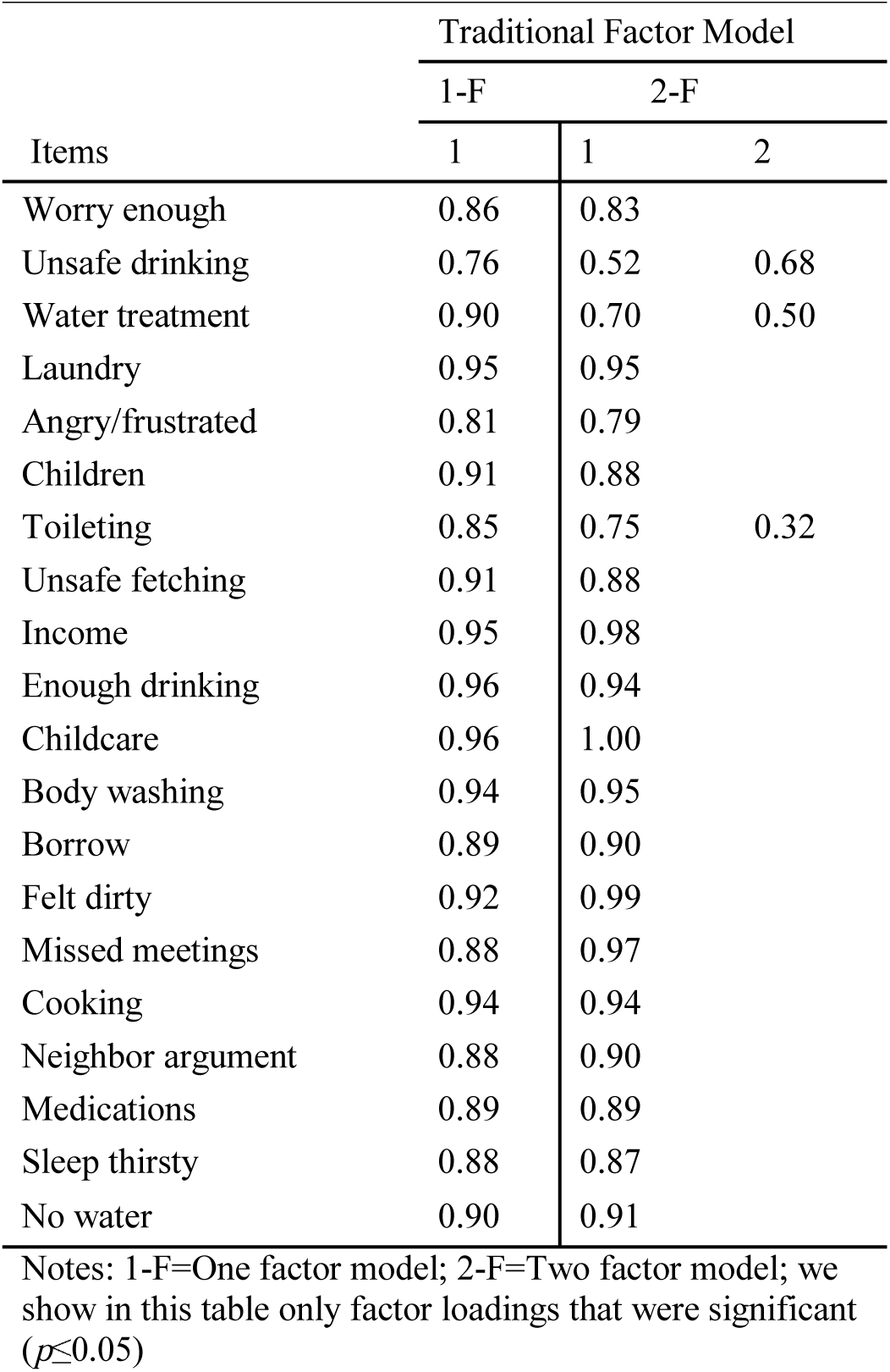
Factor loadings based on exploratory factor analysis of 20 household water insecurity items at 15 months postpartum showing one-and-two factor solutions (n=241).

All four model fit indices used in this study showed very strong support for a single dominant factor–RMSEA (0.10), CFI (0.99), TLI (0.99), and SRMR (0.04) (**S3 Table**). Therefore, we selected a unidimensional scale with 20 items. All factor loadings for the unidimensional scale were high with a minimum of 0.64 and a maximum of 0.99 all above the recommended threshold of 0.40 (Table 6). Based on the results, we hypothesized that the remaining 20 items would represent a single construct i.e. a unidimensional scale (Table 7).

**Table 7.** Final household water insecurity scale questions validated for use among postpartum women in Nyanza, Kenya.

| For each item, the questions followed the same format “In the last 4 weeks, how frequently...” |  |
| --- | --- |
| 1 | Did you worry you would not have enough water for all of your household needs? |
| 2 | Did you feel angry or frustrated that you would not have enough water for all of your household needs? |
| 3 | Did you worry about the safety of the person getting water for your household? |
| 4 | Has the time spent fetching water prevented anyone in your household from earning money? |
| 5 | Has the time spent fetching water prevented you or anyone in your household from caring for your children? |
| 6 | Has anyone in your household asked to borrow water from other people? |
| 7 | Has there not been enough water in the household to wash clothes? |
| 8 | Have you missed meetings in your community (church, funerals, community meetings, etc.) because there wasn't enough water? |
| 9 | Have you missed meetings in your community (church, funerals, community meetings, etc.) because you lacked water to take a bath and you felt too dirty to go? |
| 10 | Have you or anyone in your household had to change what was being cooked because there wasn't enough water? |
| 11 | Did you or anyone in your household had to go without washing hands after defecating, changing diapers, or other dirty activities because you didn't have enough water? |
| 12 | Did you not have enough water to wash your children's face and hands? |
| 13 | Did you or anyone in your household have to go without washing their body because there wasn't enough water? |
| 14 | Did you or anyone in your household want to treat your water, but couldn't? By treat I mean boiling, using chemicals to treat, or other ways you make your water safe to use or drink. |
| 15 | Did you or anyone in your household actually had to drink water that you thought was unsafe? |
| 16 | Did you have problems with water that caused arguments/trouble with neighbors or others in the community? |
| 17 | Has there not been as much water to drink, as you would like for you or members of your household? |
| 18 | Have you or anyone in your household not had enough water to take medications? |
| 19 | Have you or anyone in your household gone to sleep thirsty? |
| 20 | Have you had no water whatsoever in your household? |

**Notes:** For each question, participants were asked to respond to one of the following options Never (0), Rarely (1-2 times in prior 4 weeks), Sometimes (3-10 times in prior 4 weeks), Often (11-20 times in prior 4 weeks), Always (above 20 times in prior 4 weeks). Questions were asked from the least to the most severe manifestations of water insecurity.

### Tests of Dimensionality

We then tested this hypothesis using confirmatory factor model and a bifactor model on the second sample collected at 18 months postpartum. The confirmatory factor analysis test of dimensionality found partial support for our unidimensional model through the model fit indices [RMSEA (0.17), CFI (0.96), TLI (0.96), WRMR (2.05)] (Table 5). The standardized estimates from the confirmatory factor analysis were all significant at p<0.001 (Fig 2A). The bifactor model focused on accounting for unrecognized distortions created by the three items with cross loadings (Fig 2B). Reise et al. [85] suggest that where the factor loadings of the general/dominating factor are greater than the subfactor, a unidimensional factor is implied. In this analysis, the factor loadings in the dominating factor were greater than the group factor, thus pointing to a unidimensional factor. The standardized estimates on the general factor were all significant at p<0.001 (Fig 2B). Additionally, all four model fit indices [RMSEA (0.09), CFI (0.99), TLI (0.98), WRMR (0.94)] showed strong support and suggested the unidimensional hypothesis was plausible. Based on the results, we failed to reject the hypothesis that the HHWI scale consisting of 20 items was homogeneous.

**Fig 2A.**
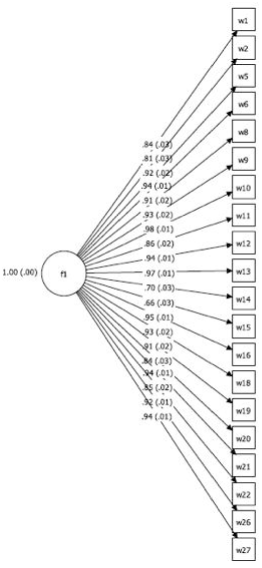
Confirmatory factor analysis with standardized estimates for household water insecurity scale at 18 months postpartum (n=186). Fig 2B. Bi-factor analysis with standardized estimates for household water insecurity scale at 18 months postpartum (n=186).

Once dimensionality was confirmed, we then summed the responses from the 20-item HHWI scale at both 15 and 18 months postpartum to create two composite scores. At 15 months postpartum, the mean of HHWI was 9.5 ± 12.2 (Mean ± SD), with a range of 0-59 (Fig 3). At 18 months postpartum, the mean of HHWI was 10.1 ± 12.4 (Mean±SD), with a range of 0-57.

**Fig 3.**
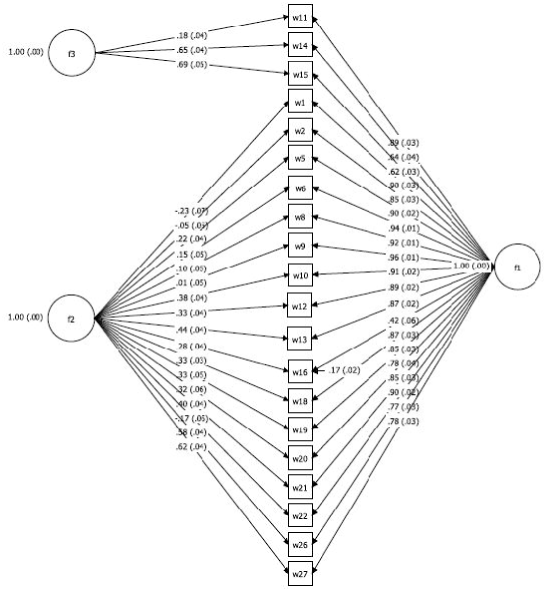
The distribution of household water insecurity scores at 15 months postpartum among women in western Kenya (n=241).

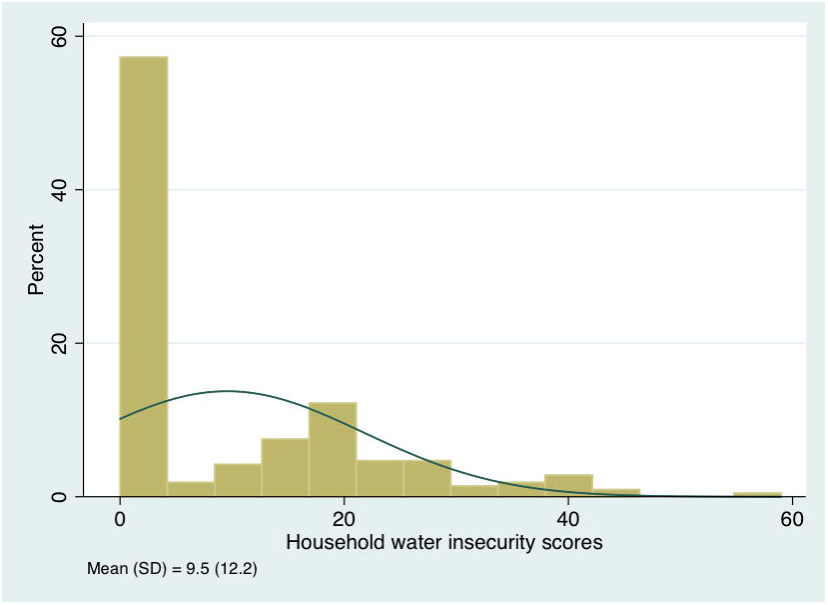

### Scale Reliability

Our test of intra-respondent reliability of the scale questions produced a strong correlation coefficient (*r*=0.76, 95% CI: 0.50-0.72; p≤0.001) between the daily responses of participants and the retrospective responses of participants on an earlier 20-item version of the scale.

In exploring the reliability of HHWI scale across time, we first computed the Cronbach’s alpha for scale items at 15 and 18 months postpartum. At both 15 and 18 months postpartum, the 20-item scale produced a Cronbach’s alpha of 0.97 (Table 8). These results suggest the internal consistency of the items at the two-time points were above all published thresholds for satisfactory reliability (α=0.70, α=90, α=95) [88,90,96]. Secondly, we assessed test-retest reliability, which correlates the scores on a given scale at two different time points to give us the coefficient of stability. Our estimation with the 20-item scale produced a positive correlation coefficient (*r*=0.62, 95% CI: 0.50-0.72; p≤0.001); however, it was below the recommended threshold (≤0.70) for the true reliability of the scale items [63,88,96].

**Table 8.** Reliability coefficients of the household water insecurity scale.

|  | Cronbach's alpha | Coefficient of Stability |
| --- | --- | --- |
| 15 months (n=241) | 0.97 | 0.62 |
| 18 months (n=186) | 0.97 |  |

### Scale Validity

To assess predictive criterion validity, we regressed maternal stress and food insecurity on HHWI score, and found HHWI to be significantly positively correlated with increased maternal perceived stress (*b*=0.12, 95% CI: 0.07-0.16, p≤0.001; β=0.34) and food insecurity (*b*=0.08, 95% CI: 0.08-0.66, p≤ 0.01; β=0.18).

To test convergent validity, our analyses showed statistically significant correlations between HHWI and total time spent per week among all households to acquire water (*r*=0.41, 95% CI: 0.23-0.57, p ≤ 0.001) and total amount of money spent on water in the last month (*r*=0.25, 95% CI: 0.12-0.37, p ≤ 0.001) at 15 months postpartum. Similar results were obtained at 18 months postpartum for total time per week acquiring water (*r*=0.38, 95% CI: 0.24-0.51, p ≤ 0.001) and amount of money spent on water (*r*=0.20, 95% CI: 0.05-0.35, p ≤ 0.01).

To assess discriminant validity, we tested if there would be a low or no relationship between HHWI and per capita household water use. This relationship was not statistically significant (*r*=0.12. 95% CI, −0.30-0.50, *p*=0.59).

We also examined the differences between ‘known groups’ on HHWI scores. Our results showed that although the magnitude of the means for the groups measured was in the expected direction, they were not statistically significant. The mean HHWI scores for participants with *E.coli* present in their household drinking water was higher (18.40 vs. 12.85; *t*=1.05, *p*=0.30; Point–Biserial *r=*0.22, Cohen’s *d=*0.44). Mean HHWI scores were higher in the dry season than in the wet season (11.15 vs. 8.69; *t*=1.43, *p*=0.12; Point-Biserial *r=*0.10, Cohen’s *d*=0.21). HIV positive women had higher mean HHWI scores than HIV negative women (11.04 vs. 8.03; *t*=-1.80, *p*=0.07; Point-Biserial *r=*-0.12, Cohen’s *d*= −0.25), as did households relying on unimproved water sources (11.67 vs. 8.45; *t*=-1.62, p=0.11; Point-Biserial *r=*-0.12, Cohen’s *d*= - 0.25) on HHWI scores.

## Discussion

A suite of rigorous qualitative and quantitative methods has yielded a 20-item scale that is valid and reliable for the assessment of HHWI among postpartum women in Nyanza region (Table 7).

Our final scale is composed of items measuring different aspects of water insecurity, yet its latent structure reflects the central assumption of unidimensionality (Tables 5 & 6). This unidimensionality is consistent with the structure of household water insecurity scales developed in Ethiopia [2,40], Bolivia [36] and Uganda [3], but differs from the work in Texas [37] and Nepal [39] where the structure of HHWI is portrayed as multidimensional. Scale dimensionality was not assessed in any of these studies with the statistical rigor used here; we encourage future studies to draw from the methods outlined here for comparable assessment of dimensionality.

The HHWI scale performed well in terms of recall bias, with a correlation coefficient of 0.76 between retrospective and prospective responses. As for reliability across time, this scale had a Cronbach’s alpha of 0.97 at 15 and 18 months postpartum, which is well above the recommended threshold for assessing the internal consistency of scales [88,90,96] and higher than other coefficient alphas reported in HHWI scales elsewhere [2,39,40]. Further, the coefficient of stability (r=0.62) attests to the strength of our scale over time. With the exception of Stevenson et al. [41] who report on a pre-post intervention study using repeated measures of household water insecurity, HHWI scales to date have not included repeated measures, which makes it impossible to compare our test-retest results to other existing HHWI scales [1].

Validity was supported in a number of ways. HHWI was positively associated with food insecurity and maternal stress, indicating predictive validity. This finding also affirms the fact that water insecurity is inextricably linked with food insecurity and has significant implications for sustainable development and poverty reduction [97,98]. The positive correlation between water insecurity and maternal stress also points to the psychosocial effects that water insecurity could have on households [4,20,99]. Future research will benefit from exploring the joint influences of food and water insecurity on health and well-being.

HHWI scores were positively associated with time spent collecting water and the amount of money spent on water, which suggests convergent validity. Water insecure households are more likely to spend more resources (time and money) obtaining water, which may lead to increased economic burden and a disproportionate burden on women who are primarily responsible for water collection [2,100]. The consequence of HHWI on women’s economic burden should be quantified in future research.

We assessed discriminant validity by examining the relationship between HHWI scores and per capita household water usage. We expected that HHWI would be different from household water use, such that there would be no relationship, and indeed, we found no statistically significant relationship between the two factors. This was consistent with the findings of Hadley and Wutich [36], Stevenson et al. [2] and Tsai et al. [3] who also found no relationship between HHWI and household water use.

As a final test of validity, we evaluated the ability of our scale to differentiate between four ‘known groups’, i.e, groups we expected to have greater HHWI: respondents with *E. coli*-contaminated drinking water vs. those without; households interviewed in the dry season vs. in the rainy season; households in which the mother was HIV-infected vs. -uninfected; and households who used improved vs. unimproved sources of water. While none of the between-group differences were statistically significant, they were all in the expected direction. We anticipate that with larger samples, we might find significant associations in the expected direction, as Tsai et al. found in Uganda [3].

Lastly, it is clear that many of the experiences of HHWI were psychosocial in nature (Figure 1). The proportion of persons who endorsed severe expressions of HHWI is of real public health relevance; 23.8% slept thirsty and 19.2% had no water in the household whatsoever at least once in four weeks (Table 4). Nearly half (46.3%) of respondents reported worrying about not having enough water and 38.3% reported feeling angry or frustrated about insufficient water in the last four weeks (Table 4). The frequency of these experiences suggests that the mitigation of HHWI in this region must be a priority for stakeholders and policy makers.

Although this study is amongst the most rigorous endeavors to develop a HHWI scale, there are a few limitations worth noting. First, the scale was developed in western Kenya; it is unlikely to be suitable for explorations of HHWI elsewhere without significant adaptation. The development of a cross-culturally validated HHWI scale is needed to assess and compare patterns of HHWI.

Second, the scale was developed among Kenyan women with young children; hence, some questions may not be applicable to households in which there are no children, e.g. Q5, Q12. Women were targeted as respondents because they are primarily responsible for water acquisition in this region [35]. We posit that this scale will be appropriate for measuring HHWI when men are primarily responsible for water acquisition and use. However, this needs to be empirically tested.

A further shortcoming is that twenty questions may be too many for some settings. The development of a more concise version of the scale, analogous to the Household Hunger Scale [101] might facilitate the implementation of this scale in diverse settings.

The study would have been strengthened if the sample used for the confirmatory factor analysis would have been from a different population entirely. Having the same participants at 15 and 18 months postpartum from the same cohort may increase common method variance and contribute to the interrelationship we find between the two time points [102]. It will be interesting to confirm the hypothetical model on a new sample both in Kenya and elsewhere to ascertain if they have the same meanings, latent factor, and factor loadings.

Lastly, our coefficient of stability, which was below the threshold of reliability (Table 8), could reflect measurement error or can be attributed to the changing conditions of participants as a result of differences in seasons or people’s living situations.

These limitations notwithstanding, this scale will permit the creation of an indicator, household water insecurity, that can be used to answer a number of questions with clinical, programmatic, and policy implications. In much the same way that our ability to measure household-level food insecurity transformed our understanding of a range of clinical outcomes, including depression [99], obesity [103], and HIV acquisition and disease progression[104], we expect this scale to be useful for understanding how water insecurity impacts nutrition, disease, and psycho-social well-being. It can also shed light on the roles that water insecurity may play in food insecurity. Further, from a policy perspective, it can be used to identify and target resources to individuals and or areas where water insecurity is highest. Finally, it can be used to assess if technological, infrastructure, or policy interventions related to water security have measurable impact.

## Conclusions

In sum, our 20-item HHWI scale (Table 7) is a reliable and well-validated measure of HHWI among women in Nyanza, Kenya. The implementation of this scale will make it possible to understand and quantify both the multi-factorial causes and consequences of HHWI (physical, mental, economic, social and nutritional). Also, the use of the scale will enable the monitoring of changes in HHWI over time and facilitate the provision of interventions to targeted households in need of support to increase household water security.

## Supporting Information

**S1 Fig.** Integration of Delphi Method with Focus Group Discussions

**S2 Fig.** Scree plot showing cut-off point for retained scale factors using exploratory factor analysis (Geomin Oblique Rotation)

**S3 Fig.** Scree plot showing cut-off point for retained scale factors using parallel analysis

**S1 Table** Delphi questions on household water insecurity

**S2 Table** Initial and final household water insecurity scale items for use among postpartum women in Kenya

**S3 Table** Description of model fit indices and thresholds used in evaluating scale development results

## Acknowledgements

We are grateful to the dedicated mothers and babies of the Pith Moromo and Pii En Ngima Study cohort, whose participation has given us a new insight to household water insecurity. A thousand thanks to our study nurses: Joy China, Joyce Bonke, and Tobias Odwar; and also to our study trackers: Benter Ogwana, Teresa Owade, and Sarah Obaje. We also wish to thank Drs. Ruth Richardson and Yolanda Brooks of Cornell University for their work on evaluating the quality of water for the presence of *E.coli*. We are grateful to Dr. Mallory Johnson for his contribution to this study. We appreciate the immense support provided by the Atkinson Centre for Sustainable Future. The content of this paper is solely the responsibility of the authors.

Contributions
The authors’ responsibilities were as follows: GOB and SLY: wrote the introduction and had primary responsibility for the final content of the manuscript; GOB: designed and performed data cleaning, analysis, wrote the methods, results, discussion, and conclusion parts of the manuscript. TBN: supervised the analysis; SLY, SC, PM, MO and PW were involved in study coordination, data collection and data management. SLY designed the study; all authors contributed to the intellectual content of the manuscript, provided critical review of the manuscript, and approved the final manuscript.

